# DRDOCK: A Drug Repurposing platform integrating automated docking, simulations and a log-odds-based drug ranking scheme

**DOI:** 10.1101/2021.01.31.429052

**Authors:** Kun-Lin Tsai, Sui-Yuan Chang, Lee-Wei Yang

## Abstract

**Motivation:** Drug repurposing, where drugs originally approved to treat a disease are reused to treat other diseases, has received escalating attention especially in pandemic years. Structure-based drug design, integrating small molecular docking, molecular dynamic (MD) simulations and AI, has demonstrated its evidenced importance in streamlining new drug development as well as drug repurposing. To perform a sophisticated and fully automated drug screening using all the FDA drugs, intricate programming, accurate drug ranking methods and friendly user interface are very much needed.

**Results:** Here we introduce a new web server, DRDOCK, **D**rug **R**epurposing **DO**cking with **C**onformation-sampling and pose re-ran**K**ing - refined by MD and statistical models, which integrates small molecular docking and molecular dynamic (MD) simulations for automatic drug screening of 2016 FDA-approved drugs over a user-submitted single-chained target protein. The drugs are ranked by a novel drug-ranking scheme using log-odds (LOD) scores, derived from feature distributions of true binders and decoys. Users can submit a selection of LOD-ranked poses for further MD-based binding affinity evaluation. We demonstrated that our platform can indeed recover one of the substrates for nsp16, a cap ribose 2′-O methyltransferase, and recommends that fluralaner, tegaserod and fenoterol could be repurposed for the COVID19 treatment with the latter two being confirmed in SARS-CoV2 suppression experiments. All the sampled docking poses and trajectories can be 3D-viewed and played via our web interface. This platform shall be easy-to-use for general scientists and medicinal researchers to carry out drug repurposing within a couple of days which should add value to our timely responses to, particularly, emergent disease outbreaks.

**Availability and implementation:** DRDOCK can be freely accessed from https://dyn.life.nthu.edu.tw/drdock/. (Due to the hardware upgrade, the service is NOT available before 7/18, 2021)

## 1 Introduction

In emergent diseases, re-use or recombination of old drugs, known as drug repurposing, can always be the first available remedy to respond to the diseases for their evidenced safety and preferred bioavailability (Ashburn et al., 2004; Barratt et al., 2012; Jin et al., 2014; Liu et al., 2018; Oliveira et al., 2018). In fact, statistics showed there are less and less new molecule entities (NMEs) (Avorn, 2015; Cleary et al., 2018; Paul et al., 2010) entering the market with an approval from FDA in the past decade for their inevitable side effects, unexpected low efficacy, long developmental time and accompanied high cost (Hughes et al., 2011; Mohs et al., 2017; Vohora et al., 2018).

Identification of drugs that could be repurposed for new targets requires an efficient and accurate high throughput drug screening and repurposing platforms, where the former uses all the available chemical leads, easily millions of them, and the latter uses FDA-approved drugs, or those that are still in the clinical trials (Ekins et al., 2007; March-Vila et al., 2017). One common practice uses small molecular docking (Forli et al., 2016; Gilson et al., 2007; Hsu et al., 2011; Kontoyianni M., 2017; Ma et al., 2011; Murgueitio et al., 2012), where different binding positions/orientations and configurations of a drug (termed “poses”) are sampled in a protein and the resulting poses are rank-ordered by the docking scores or software-defined binding enthalpy. These docking simulations are performed in a vacuum with no dynamics involved, and, depending on the scoring functions, limited ligand entropy (Chang et al., 2007) change upon binding is considered.

The intrinsic protein dynamics (Eyal et al., 2015; Li et al., 2017; Yang et al., 2005; Yang et al., 2006; Yang et al., 2009; Zhang et al., 2020) has been proved crucial to modulation of ligand binding (Li et al., 2014; Yang et al., 2005) and used to assess the druggability of a protein target (Bakan et al., 2012; Lee et al., 2019). When excited by specific physicochemical perturbations, a small set of these intrinsic normal modes are recombined to drive protein conformational changes near the equilibrium state (Bakan et al., 2009; Yang et al., 2014). Given the modern computing power, molecular dynamics (MD) simulations have been largely used to describe the dynamic behavior of drugs-target interactions. The simulation trajectories of selected docking poses with the highest affinities (Alonso et al., 2006; Chang et al., 2016; Guterres et al., 2020) can be further analyzed for binding free energy evaluation at body temperature and room pressure, with neutralizing ions solvated in the explicit solvent (Guterres et al., 2020; Salmaso et al., 2018; Takemura et al., 2013).

On the other hand, most of the docking scoring functions are calibrated using a set of high qualities protein-ligand complex structures annotated by experimentally assayed binding affinities (Ballester et al., 2010; Cao et al., 2014; Huey et al., 2007; Koes et al., 2013; Trott et al., 2009; Wang et al., 2002; Wang et al., 2011;). Due to the nature of structural biology, those experimentally “seen” in x-ray crystallography are usually stable co-crystallizer-protein complexes with relatively high affinity. Therefore, the scoring functions derived from these high-affinity ligands may fail to deprioritize low-affinity ligands that might only form transient binding with the target proteins since they are not included in the calibration process. This could introduce false positives of low-affinity drugs that are inappropriately prioritized in the docking process. If aforementioned simulation-based refinement and re-ranking are to be applied for only top-ranked drugs from the docking procedure, these wrongly prioritized low-affinity ones would result in a waste of computing resources. To best utilizing the computing resources and promote the success rate of drug repurposing, an accurate drug ranking method that is capable of distinguishing the effective drugs (true binders) from transient binders (decoys) is required.

In the past decades, there were web servers reported to provide small molecular docking and compound screening for drug discovery (Singh et al., 2020). SwissDock (Grosdidier et al., 2011), a reputed pioneer of docking web server, provides online computing resources for one-target-one-ligand docking service with limited acceptance for batched ligands-submission. iScreen (Tsai et al., 2011) provides cloud-computing resources for screening more than 20,000 traditional Chinese medicines (TCM) against a user-provided target by the docking program PLANTS (Korb et al., 2007; Korb et al., 2009). DockCoV2 (Chen et al., 2020), a target-centric drug repurposing database in response to the COVID-19 pandemic, deposits freely accessible simulated docking poses with no further MD-refinement for ∼3000 administration-approved drugs against seven focused protein targets in SARS-CoV-2. Inverse docking web services, such as ACID (Wang et al., 2019) and idTarget (Wang et al., 2012), achieve drug repurposing by reverse screening, one drug against many drug targets, to fish promising targets to which the drug could bind. However, it does not allow to repurpose old drugs for newly emerged drug targets. ACFIS (Hao et al., 2016), another web service for *de novo* drug design, applies a fragment-based strategy by adding 2883 small fragments decomposed from compounds database, including FDA-approved drugs, to a core fragment derived from the input ligand in the user-provided protein-ligand complexed. This generates a set of new compounds with increased binding affinities to the target protein evaluated based on a few picosecond MD simulations. Despite these marked progress in making available customized online drug-screening services, there has not been a fully automated functional web server, allowing any uploaded PDB structure that can also be optionally relaxed by MD as a drug target over which several thousand of FDA-approved drugs are docked, followed by ≥10 nanosecond explicit-solvent MD simulations and trajectory-based binding affinity evaluation. To be responsive to emergent infectious diseases and diseases without gold standard therapeutics, the aforementioned online service can be helpful to assess the binding affinity of all the FDA-approved drugs with essential genes in emergent pathogens, considering protein-drug dynamics in water at body temperature.

Here, we introduce DRDOCK, **D**rug **R**epurposing **DO**cking with Conformation-sampling and pose re-ran**K**ing, a new web server that integrates molecular docking and MD simulations for automatic virtual drug screening for drug repurposing of 2016 FDA-approved drugs. To improve the ranking accuracy of these drugs, we introduced a benchmark comprising drug-target pairs and an FDA-drug-specific metric to evaluate the goodness of any ranking method. With those, we developed two ranking methods, where in one of them, we developed the logarithm of odds (LOD) scores derived from pose features statistics (namely, pose affinity, distance to the binding pocket and entropic effect of poses) from true binders and decoys. Our web service implemented the LOD score, MD-based conformational sampling and MM/GBSA calculations to improve the screening accuracy in a relatively safe chemical space and therefore accelerate the drug repurposing.

## 2 Materials and methods

### 2.1 *In silico* FDA-approved drug library

2016 FDA approved drugs are collected and combined from the catalogs of MedChemExpress (MCE) FDA-Approved Drug Library (Cat. No.: HY-L022) and Enzo Life Sciences, Inc. Screen-Well^®^ FDA Approved Drug Library (Version 1.5). The model 3D structures of drugs are built using BIOVIA Discovery Studio (Dassault Systèmes BIOVIA, Discovery Studio Modeling Environment, Release 2017, San Diego: Dassault Systèmes, 2016) with the protonation states of ionizable groups assigned at pH = 7. The input PDBQT files of drugs for docking are generated using AutoDock Tools (Morris et al., 2009), and the parameters required for MD simulations are assigned by Antechamber (Wang et al., 2006) based on GAFF2 (Wang et al., 2004) force field with am1-bcc (Jakalian et al., 2002) method for charges calculation.

### 2.2 Protein preprocessing

DRDOCK is designed for drug screening on a fully patched, single-chain protein target in the PDB format. The user-uploaded protein target is automatically processed and prepared for docking and MD simulations, including the prediction of protonation states of Ionizable groups by using PROPKA3.1 (Olsson et al., 2011; Søndergaard et al., 2011), the flipping of the side-chain orientation of asparagine (Asn), glutamine (Gln), and histidine (His) for optimized hydrogen-bonding and the avoidance of NH2 clashes with surrounding toms adjusted by Reduce (Word et al., 1999), and automatic disulfide bonds detection (S-S distance < 2.5 Å). The input PDBQT file for docking is generated using AutoDockTools (Morris et al., 2009), and the ff14SB force field (Maier et al., 2015) is used for MD simulations.

### 2.3 Benchmark set curated for the development of drug ranking methods

To develop the drug ranking methods and compare their performance, the complex structures for a set of known target proteins and FDA-approved drugs were extracted from the Protein Data Bank (PDB) (rcsb.org; Berman et al., 2000). We first collected 48 chemical IDs of FDA drugs that were newly approved during 2010-2016 gathered in the article published by Westbrook et al (2019) (the drugs reported to target the same target proteins were excluded for simplicity). Among these 48 approved drugs, 44 were included in our 2016 FDA-approved drug library. We then fetched the PDB IDs in which the structures were co-crystallized with one of the 44 drugs, eventually arrived in 20 drug-target complexes (see Supporting Info for details), listed in Table 1. We called the FDA-approved drug in the complex structure as “true binder”.

**Table 1.**
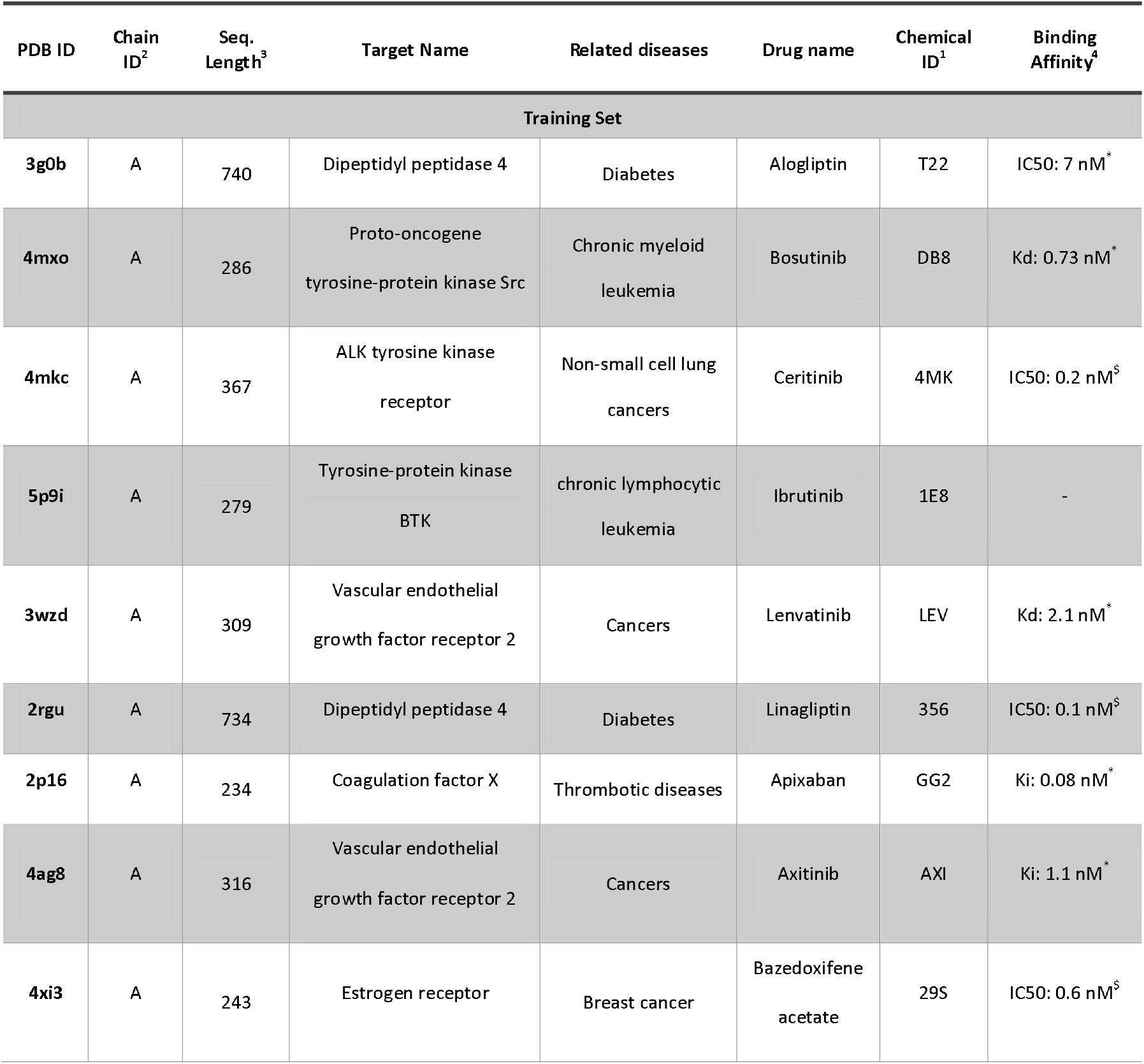
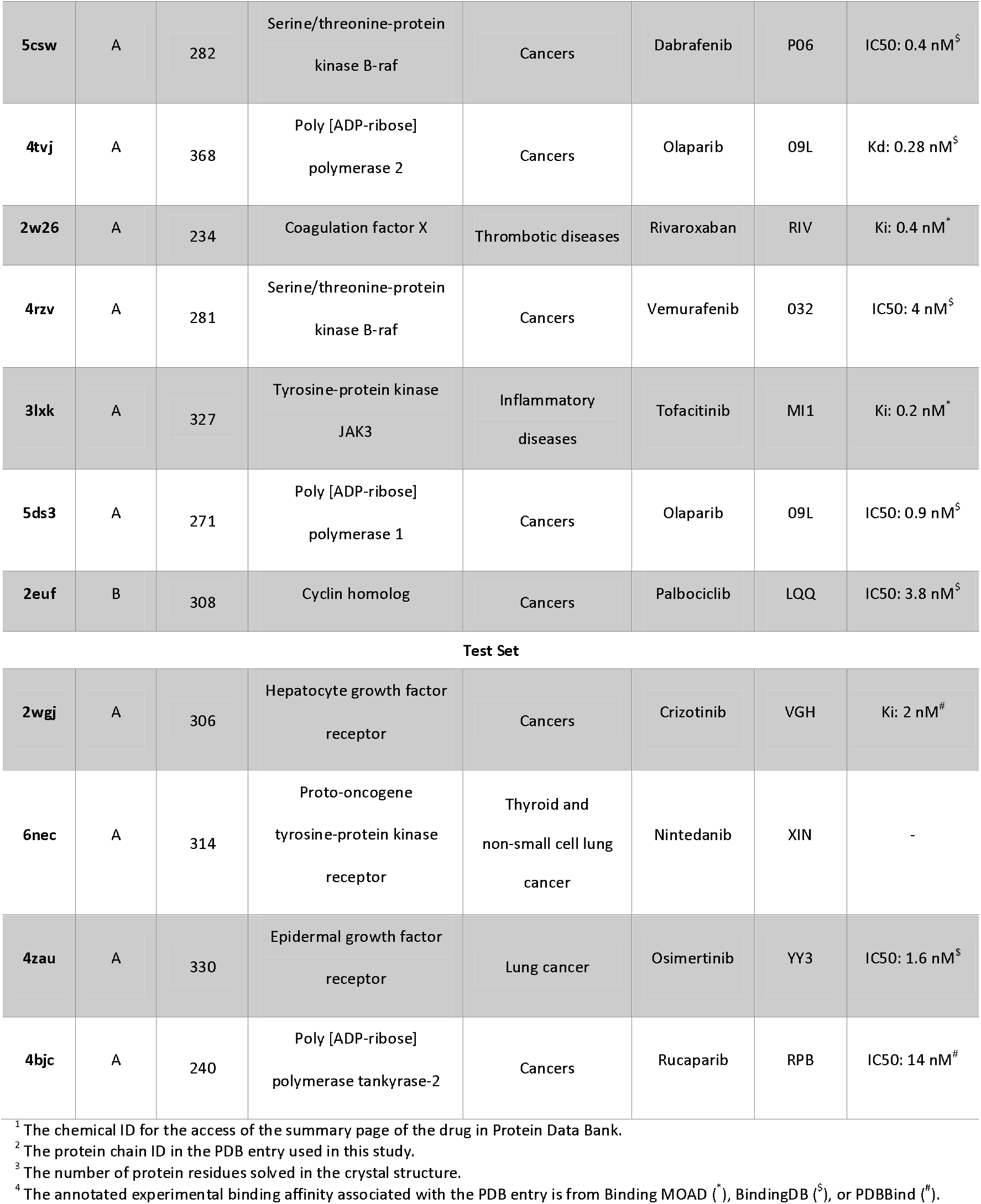
The collected 20 PDB complexes containing FDA-approved drugs used in this study.

To generate the docking poses and their associated pose features (see the next section), each of the 20 complex structures (Table 1) was docked with our in-house built 2016 FDA-approved drugs by Autodock Vina (Trott et al., 2009). We defined the true binding pose of the true binder as the docking pose that among all has the shortest distance (measured from its mass center) to the mass center of the true binder’s heavy atoms in their co-crystallized positions. The decoys (Graves et al., 2006) are defined as all the sampled poses of the 2016 FDA-approved drugs that are not the true binding pose.

### 2.4 Virtual drug screening and its resulting features

AutoDock Vina (Trott et al., 2009) was used for drug screening, where each of the 2016 FDA-approved drugs was docked into the target protein and allowed sampling for 20 poses. The docking box was set to cover the whole structure of the target protein with an additional 5 Å margin patched on each side of the docking box. The exhaustiveness was set to 50 to give accurate screening results within a reasonable time (∼16hrs for the 2000+ drugs using 120 Intel Xeon E5606 CPU cores). The docking results were analyzed based on (A) the predicted binding affinity by AutoDock Vina, (B) the distance between the docking pose to the assigned target residues, and (C) the size of the poses cluster, as detailed below.

For Vina-predicted binding affinity, we used the original value (affinity) or the value divided by the square root of the number of drug heavy atoms (normalized affinity). The distance was measured as the closest distance between any heavy atoms of the drug pose to the center of mass (COM) of the assigned functional residues in a protein target (termed as “closest distance”), or the distance between COMs of a drug pose and assigned target residues (termed as “COM distance”). We also grouped similarly oriented poses into clusters. Clustering the docking poses into groups is to approximate the entropy contribution by the size of the clusters, where a large cluster suggested a less entropy loss during the drug binding process. To cluster the docking poses of each drug, we used the hierarchical clustering (Sokal and Michener, 1958) with a cutoff of 4 Å. Two types of distances were used - the distance between COMS of two poses (termed as “COM-based clustering”) and the “adjacency” of docking poses (termed as “adjacency-based clustering”, see below). In adjacency-based clustering, poses were clustered based on whether any of their heavy atoms come within 1 Å. The contact map of 20 poses × 20 poses was then used to identify the connected components, the “pose clusters” here, using an improved version of Tarjan’s algorithm (Pearce, D.J., 2005) implemented in SciPy library (Virtanen et al., 2020). The three drug ranking algorithms introduced in the next section differ in their ways to integrate these aforementioned features, with an aim to prioritize the true binding pose/drug as front as possible among the 2015 competitors against the true binder’s corresponding protein target.

### 2.5 *Ad hoc* method for drug ranking

The *ad hoc* ranking method is adapted and modified from our previous work (Liu et al., 2018), which is composed of two stages hierarchical classifying and sorting based on aforementioned three features (see Figure S1 for details) except that the cluster size here refers to the number of poses in a cluster that rank top 10 in affinity (out of 20) for that drug. In the first stage, a representative pose was chosen among the sampled 20 poses for each drug. This was achieved by first applying a binary classification to group and prioritize those poses that were within 5 Å of the target site. Within each of the resulting two groups, a second binary classification was applied to each group where we prioritized the poses that belong to the largest clusters formed by the sampled 20 poses. Within each of the resulting four groups, the poses were then ordered based on the predicted binding affinity. The first pose in the top-ranked group was then chosen as the representative pose for that drug. The selected 2016 representative poses for 2016 drugs from the first stage were then used to determine the ranking of the drugs in the second stage. The drugs were again first binary-classified to prioritize the drugs that were within 5 Å of the target site. The drugs in each of the two groups were then ranked based on the predicted binding affinities.

### 2.6 Log-odds score (LOD score)

To develop the LOD score, the ranking of drugs is decided by a score based on the distance of pose features’ statistics distributions, sampled within our benchmark set, from the true binders and the decoys. To get such statistics, we pooled all the generated poses from the 16 protein-drug complexes in the training set and derived the distribution of the true binding poses and decoys for each of the features associated with each pose. These distributions provided the statistical basis to annotate a sampled docking pose to be a true binder or decoy, based on its relative probability. Specifically, we calculated a log-odds (LOD) score for each of the sampled poses according to the following equation (Eq. 1):

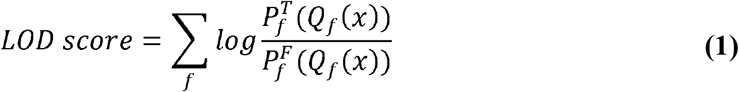

where *x* represents a sampled docking pose; *f*represents one of the specified features we mentioned in section 2.4, e.g. the pose affinity, the pose’s distance to the target site, and the size of poses cluster. *Q*_*f*_ (*x*) is the value of feature *f*for the pose *x* is located. 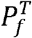 and 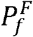 are the probability distributions of feature *f*for docking poses sampled from the true binders and the decoys, respectively. Noted that *Q*_*f*_ (*x*) are binned values and therefore different *Q*_*f*_ (*x*) values within the same bin have the same 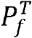 or 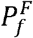 values. If there is no count for a bin, a small probability (10^−7^) is assigned for that bin. The LOD score takes the idea of relative entropy, representing how more or less likely a given sampled pose can result from a true binder than from a decoy. A higher LOD score means the pose is more likely from a true binder. The 2016 drugs are rank-ordered based on their best LOD score among the 20 sampled poses.

### 2.7 Feature ranking score (FRS)

In the FRS method, the values *for a single feature* to all the poses of the 2016 drugs are sorted. A lower drug affinity and drugs’ distances to target sites, or a larger size of pose clusters score correspond to a higher rank, starting from one. For each pose, the ranking for each of the three features is summed to obtain an FRS score such that

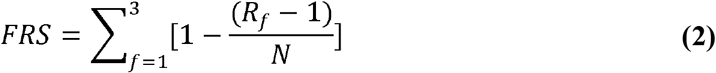

where *f* represents one of the three features. *R*_*f*_ represents the ranking of a pose for the feature *f*. *N* is the total number of the sampled poses from all the 2016 drugs. The pose with the highest FRS in each drug is used as the representative pose for that drug, which is then sorted to obtain the final ranking for the 2016 drugs.

In the weighted FRS, the scores from different features are linearly combined such that

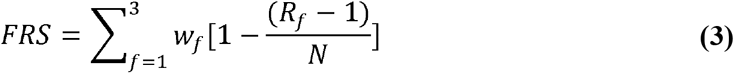

where *w*_*f*_ is the corresponding weight for the feature *f*.

To determine the weights associated with each feature, we constrained the range of values of the weights such that 0 < *w*_*f*_ < 1 *and* ∑_*f*_ *w*_*f*_ = 1.

We then sought the set of {*w*_*f*_} that minimized the following average rank

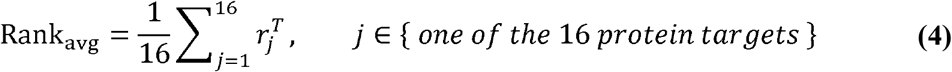

where 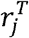 is the rank of the true binder in *j*th target protein of the training dataset. To search for the optimal value of {*w*_*f*_}, we applied two search methods: grid search and sampling search. Let the weights of the three features be *w*_1_, *w*_2_, and *w*_3_. In the grid search method, we allowed two of the weights, said *w*_1_ and *w*_2_, to be any value in the set {0, 0.1, 0.2, …, 0.9, 1}, and exhaustively searched the set of (*w*_1_, *w*_2_, 1-*w*_1_-*w*_2_) for the combination that minimized the loss function (Eq. 4). In the sampling search method (*Weisstein, MathWorld*), we sampled (*w*_1_, *w*_2_, *w*_3_) according to the following equation (Eq. 5).

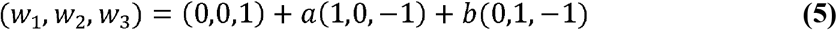

where *a* and *b* were sampled uniformly from 0 to 1, and we kept the set (*w*_1_, *w*_2_, *w*_3_) where *w*_3_>= 0. We sampled 10,000 sets of such (*w*_1_, *w*_2_, *w*_3_) and chose the one that gave the smallest loss (Eq. 4).

### 2.8 Performance evaluation of the ranking methods

To use the testing set to evaluate a certain ranking method first parametrized over the training set, the total number “16” in Eq. (4) is replaced to “4” for the four drug-protein pairs in the test set (Table 1). We also defined the success rate of the true binders as the ratio of the training or test set being in the top 100 (∼5%) or top 200 (∼10%) among all the examined drugs.

### 2.9 Molecular dynamics (MD) simulations and MM/GBSA

While docking serves as a fast screening tool for *in silico* drug screening, as a trade-off, the lack of considering the solvation effect and molecular dynamics involved in the protein-ligand interaction introduces false positives (Pantsar et al., 2018). Explicit solvent MD simulations using classical force fields optimized for biological molecules are recognized to better describe the protein-ligand binding dynamics in a timescale allowed for weak binders to dissociate (Huang and Caflisch, 2011; Liu and Kokubo, 2017; Liu et al., 2018; Mollica et al., 2015). The resulting trajectories further allow accurate binding free energy evaluation by MM/GBSA, where the molecular mechanics of protein-ligand complexes and a continuum solvent model accounting for the solvation energy calculation are considered suitable to assess relative binding affinity for a set of drugs against the same pocket (Ahinko et al., 2018; Liu et al., 2018; Sun et al., 2014; Yang et al., 2011; Zhang et al., 2017).

The MD simulations for user-selected poses were performed using the OpenMM package (Eastman et al., 2017) in an explicit solvent. The simulation model of the protein target-drug binding pose complex was prepared using the LEaP program in AmberTools (Case et al., 2020). The complex was solvated using an explicit solvent of the TIP3P water model, with at least 10 Å of water layer patched on each side of the water box between the protein target and the box boundary. Sodium and chloride ions were used to neutralize the system to achieve a salt concentration of 100 mM. The system was first energy minimized for all the hydrogen, waters, and ions positions, leaving the remaining atoms restrained using a force constant of 10 kcal/mol/Å^2^. This was followed by the energy minimization for atoms including also protein heavy atoms, leaving the protein C-alpha atoms and drug heavy atoms restrained using a force constant of 2 kcal/mol/Å^2^. The system was then heated to 320 K in NVT ensemble for 1 ns and equilibrated for another 1 ns in NPT ensemble 310 K and 1 atm. The protein Cα atoms and drug heavy atoms were weakly restrained (0.1 kcal/mol/Å^2^) during the NVT heating and NPT equilibration. The equilibrated system was then subject to a 10 ns simulation using the same condition as in the equilibration except for the removal of all restraints, and the snapshots were sampled for every 0.1 ns, resulting in a trajectory composed of 100 frames. Langevin dynamics with a collision frequency of 2 ps^-1^ and a Monte Carlo Barostat with the volume rescaled every 25 time steps were applied for temperature and pressure controls, respectively. The particle mesh Ewald (PME) method was used to calculate the energy of non-bonded interaction with 10 Å as the distance cutoff. All the bonds involving hydrogens were constrained by SHAKE algorithm (Miyamoto and Kollman, 1992). The binding free energy approximated by the enthalpy contribution between the target protein and drugs for each sampled snapshot was calculated by MM/GBSA methods (Genheden et al., 2015; Greenidge et al., 2012; Kollman et al., 2000; Massova et al., 2000) with MMPBSA.py module (Miller et al., 2012) in the AmberTools (Case et al., 2020). The simulation results were summarized using the designed indicators, including the mean binding free energy from MM/GBSA calculation over sampled snapshots, the drug leaving time when the drug center of mass moving away from the target site more than 10 Å, and the largest distance of the drug COM to the target sites sampled during the simulations. A simulated drug got a higher rank if having a favorable mean binding free energy and staying long enough in the target site (or never leaving) during the 10 ns simulation. All the processing and analysis of the docking poses and MD trajectories were performed by ParmEd (http://parmed.github.io/ParmEd), Cpptraj (Roe et al., 2013), PyTraj (Nguyen et al., 2015), and MDAnalysis (Gowers et al., 2016; Michaud-Agrawal et al., 2011).

### 2.10 SARS-CoV2 Plaque Reduction Assay

The virus strain used in this study (hCoV-19/Taiwan/NTU13/2020; Accession ID: EPI_ISL_422415) was isolated from sputum specimen obtained from a SARS-CoV2-infected Taiwanese patient. The sample was first propagated in Vero E6 cells in DMEM supplemented with 2 μg/mL tosylsulfonyl phenylalanyl chloromethyl ketone (TPCK)-trypsin (Sigma-Aldrich)(Pathak et al., 2020). Culture supernatant was harvested when cytopathic effects (CPE) were seen in more than 70% of cells, and viral titers were determined by a plaque assay. Plaque reduction assay was performed to determine the antiviral activity of test compounds. Briefly, Vero E6 cells were seeded to the 6-well culture plate in DMEM with 10% FBS and antibiotics one day before infection. VeroE6 cells were infected by SARS-CoV-2 virus (50−100 plaque forming unit, pfu) for 1 h at 37oC. After removal of virus inoculum, the cells were washed once with PBS and overlaid with 1 mL medium containing test compounds at indicated concentration and 1% methylcellulose for 5 days at 37oC. After 5 days, the cells were fixed with 10% formalin overnight. After removal of overlay media, the cells were stained with 0.5% crystal violet and the plaques were counted. The percentage of inhibition was calculated as [1-(VD/VC)]×100%, where VD and VC refer to the virus titer in the presence and absence of the inhibitors, respectively. The minimal concentrations of compounds required to reduce 50% of plaque numbers (EC50) were calculated by regression analysis of the dose-response curves generated from plaque assays. For each data point, the measurements were repeated three times to yield the averaged number, standard deviations and standard errors.

## 3 Website

### 3.1 Architecture

DRDOCK is composed of a central web server and distributed thread pools and data storage (Figure 1A). The web server plays as an interface to serve the web pages to the client browsers and receive the user requests. Once a service query is received, the web server processes the incoming query and sends compute-intensive tasks to the internal processing units-specific (CPUs and GPUs) thread pools. The web server dispatches the tasks to the specific type of thread pool through a job dispatcher hosted on a separate Redis server (https://redis.io/). The CPU thread pool is provided by an HPC cluster made of 11 nodes, each comprised of two CPUs of Intel(R) Xeon(R) X5660 (12 cores in total), for CPU-intensive tasks, such as docking and MM/GBSA calculations. The GPU thread pool equips with four GeForce GTX 980 Ti GPU cards and is responsible for carrying out MD simulations with AMBER ff14SB force field (Maier et al., 2015). The progress of the task is then reported to the web server through the internal LAN by HTTP(S). All the web server and threads can share the same files stored on a central Network Attached Storage (NAS). The portal webpage is programmed with Django Web framework (https://www.djangoproject.com/). The user-specific information and job configuration are stored using PostgreSQL https://www.postgresql.org/).

**Figure 1.**
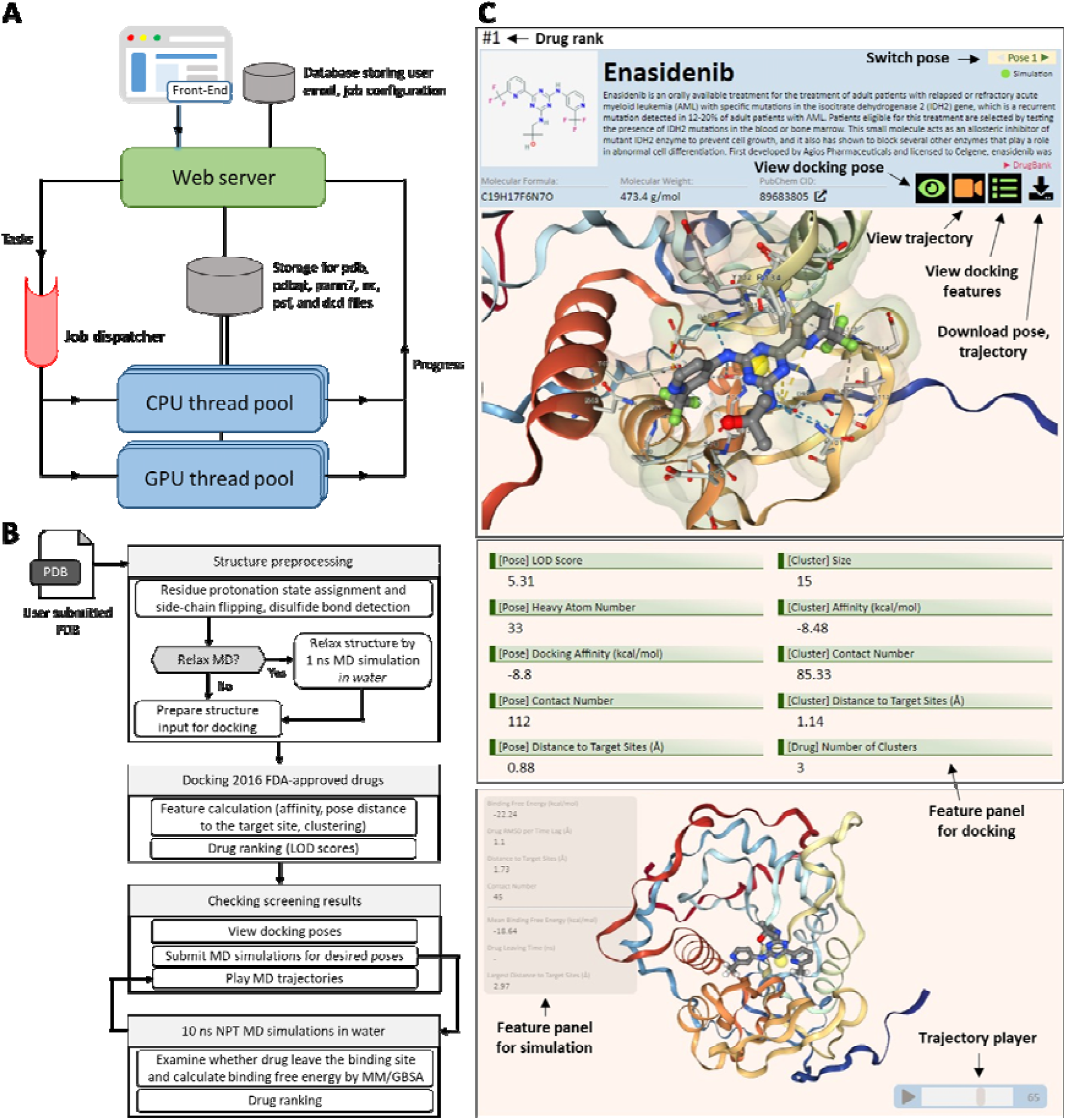
DRDOCK web service. **(A)** The architecture of DRDOCK. The central web server serves as the portal server serving the website and responds to any incoming requests. The CPU-intensive docking and GPU-intensive simulation jobs are sent from the web server separately to the corresponding thread pool on distributed machines through the job dispatcher. The progress of jobs is reported back to the web server through an internal LAN. All resulting files are kept in a central Network Attached Storage (NAS) that allows access from all the machines. A separated relational database hosted on the web server is used to save the user information and job configurations. **(B)** The workflow in DRDOCK. DRDOCK begins with preprocessing the user-submitted protein target. Users can choose to directly use the input structure for docking or optionally perform a one ns MD simulation to minimally relax the structure in water before docking. The preprocessed structure is then used to dock 2016 FDA-approved drugs, where the LOD scores calculated from the docking poses are used for drug ranking. Users then view the docking poses on the online results page and pick interesting docking poses to simulate the dynamic protein-drug interaction in 10 ns NPT MD simulations with water. The resulting trajectory is used for binding free energy calculation by MM/GBSA and can be played on the online results page. **(C)** The functionality and info-display in the drug panel. The user can visualize the docking pose by clicking the eye-shape “view docking pose” button, and an extended space showing the binding mode of protein and drug will show up (top), clicking the “view docking features” button would render us a list of properties derived from the docking results (middle). Clicking the video-shape of “view trajectory” button pops open an NGL viewer (Rose et al., 2018) and plays the MD animation for the interacting drug-target complex if the pose were chosen for simulations. With this, MM/GBSA-estimated binding free energy is listed on the side, which is important for re-ranking the docking results. The pose and trajectory can be downloaded from the “download” button.

### 3.2 Workflow

The workflow in DRDOCK can be summarized into four modules: structure preprocessing, docking/ranking 2016 FDA-approved drugs, 10 ns NPT MD simulation in water, and result rendering (Figure 1B). The user-submitted protein target in PDB format is first preprocessed before docking, including the determination of the pKa and protonation state of ionizable residues, optimizing the orientation of rotatable residue side-chains, and disulfide bond detection (Methods 2.2; see Supporting Results for more details). As an optional service, users can choose to relax the target structure (Chang et al., 2016), without bound co-crystallizers, by 1 ns MD simulation in water before the docking. The service then docks 2016 FDA-approved drugs to the target, and features inferred from docking poses (Methods 2.4) are used to calculate the LOD scores for drug ranking (Methods 2.6). Users then view the docking poses online on the results page and submit the interested docking poses for 10 ns NPT MD simulations of the protein-drug complexes in water. The resulting trajectories are used for assessing the drug binding stability (i.e., whether the drugs stay in the binding site through the simulation) and binding free energy by MM/GBSA (Methods 2.9). The trajectories are then updated to the results page for the playback of manipulatable 3D animations.

### 3.3 Result rendering

DRDOCK is designed for allowing real-time access to the latest calculation results while the entire drug screening is still in the process. The drugs and their docking poses shown in the results page are rank-ordered based on the LOD scores, described in the Methods. The drug name, 2D structure, drug properties, and external link to PubChem (Kim et al., 2019) can be viewed in each drug panel (Figure 1C, top). Docking poses for a drug can be selected by the switch at the top right (Figure 1C, top). Any docking pose can be further submitted for a 10 ns MD simulation (Figure 1C, and Figure S2B). When the simulation is finished, a superposed trajectory can be visualized in the embedded NGL viewer by clicking the “view trajectory” button (Figure 1C, bottom), and the corresponding features and the estimated binding free energies for each frame can be seen in the floated left panel (Figure 1C, bottom). For a better use of the computing resources, simulations for at most 20 selected poses can be submitted for each uploaded protein target. Finally, a complete list of rank-ordered drugs, all the docking poses, MD simulation trajectories, calculated features can be downloaded by clicking the “download” button (Figure 1C, and Figure S2B).

## 4 Results

We treat the drug screening problem as a ranking problem in order to reduce the disparity coming from different docking software used, different sampling depth and parameterization details of the drug force field etc. With this and through DRDOCK, given the consistency in an automated platform, the same drug on two different protein targets might be compared but more recommended comparison would be to examine the ranking of the drug in individual targets. Our goal here is to assess drug ranking methods based on how each method can prioritize the known true binders to the top of the 2016 FDA drugs, as high as possible.

We obtained a benchmark set of 20 drug-target complexes derived from a 2016 drug library (Westbrook et al., 2019; see Methods). We defined the co-crystallized drugs complexed with the corresponding target proteins as the true binders; for each target, there is only one true binder but 2015 decoy drugs (see also details on pose choice for the true binder). FDA drugs by design should target the best a single target of interest with the IC50 or Kd at nanomolar range and are not supposed to bind as well with other targets. For currently 2000+ FDA drugs that target nearly 900 protein targets (Santos et al., 2017), in average, more than 2 drugs “well bind” a single target. However, by how FDA drugs are designed and by the statistics where per drug targeting 0.5 protein in average, it should be reasonable to assume that the “cognate” or “native” drugs bind their designed targets the best or nearly the best, while the rest of the FDA drugs should not bind the same targets as well as the “native”, hence our seemingly bold assumption of 1 true binder and 2015 hit decoys for each target. The 20 structures in benchmark set were randomly split into 16 and 4 respectively as the training set used for method calibration and the testing set for evaluation of method performance. To get the atomic interactions of the drugs to the target proteins, each drug was docked or re-docked (for true binders) to the target proteins using a global search. At most 20 docked poses resulted for each drug and the features associated with the docking poses were calculated (Methods). We generated ∼ 20 * 2016 * 20 poses for the 20 target proteins, where each pose was associated with the aforementioned three features (Methods).

Three drug ranking methods were evaluated using our benchmark set – using the LOD score, feature ranking score (FRS) and an *ad hoc* method modified from our previous study (Liu et al., 2018). To properly defined the LOD scores for our system, we needed to first identify relevant features that can render distinct probability distributions of true binders and decoys for that feature. Then, the most distinguishing features were used to derive the LOD scores, by summing which for each feature drugs are rank-ordered and the performance of the method is evaluated.

Feature ranking score (FRS) rates a pose by its ranking pertaining to each of examined features. As described in Eqs. 2 and 3, FRS is a sum of un-weighted (Eq. 2) or weighted (Eq. 3) ranking-normalized scores such that the highest rank in a certain feature gets the unity value while the lowest rank is nearly zero. To determine the weight of each feature in the linear combination, a grid-search method and a constrained sampling method are used to minimize the average rank of the true binders. The drugs are then ranked based on the pose with the highest FRS for each drug (Methods). As a comparison, the *ad hoc* approach was proposed (Liu et al., 2018) and its description can be found in Methods.

We quantified the difference between the distributions by Kolmogorov–Smirnov (K-S) test (Pratt, J.W., 2012) and Kullback–Leibler (K-L) divergence (Cover et al., 2006). A feature is considered “preferred” if it gives a large K-S statistics or K-L divergence (the “distance” between two distributions) between the distribution of true binders and that of decoys. The two distributions for each of the features are shown in Figures 2 to 4. For the binding affinity, the distributions of the original values between the true binders and decoys (K-S statistic: 0.775, K-L divergence: 2.981) were more different than that of the normalized affinity (K-S statistic: 0.188, K-L divergence: 0.273) (Figure 2). For the distance to the target sites, the true binders had shorter COM distances compared to that of decoys (K-S statistic: 0.828, K-L divergence: 2.415), and the difference was more pronounced than the distances measured using the closest drug heavy atoms (K-S statistic: 0.557, K-L divergence: 0.595) (Figure 3). Regarding the cluster size, the clusters grouped by adjacency-based clustering had stronger power to distinguish true binders from decoys (K-S statistic: 0.274, K-L divergence: 0.458) than that grouped by COM-based clustering (K-S statistic: 0.156, K-L divergence: 0.271) (Figure 4). Therefore, we used the regular affinity value as the measurement for the pose binding affinity, the COM-distance as the measurement for the pose distance to the target sites, and the cluster size resulted from the adjacency-based clustering as the measurement of the pose cluster size.

**Figure 2.**
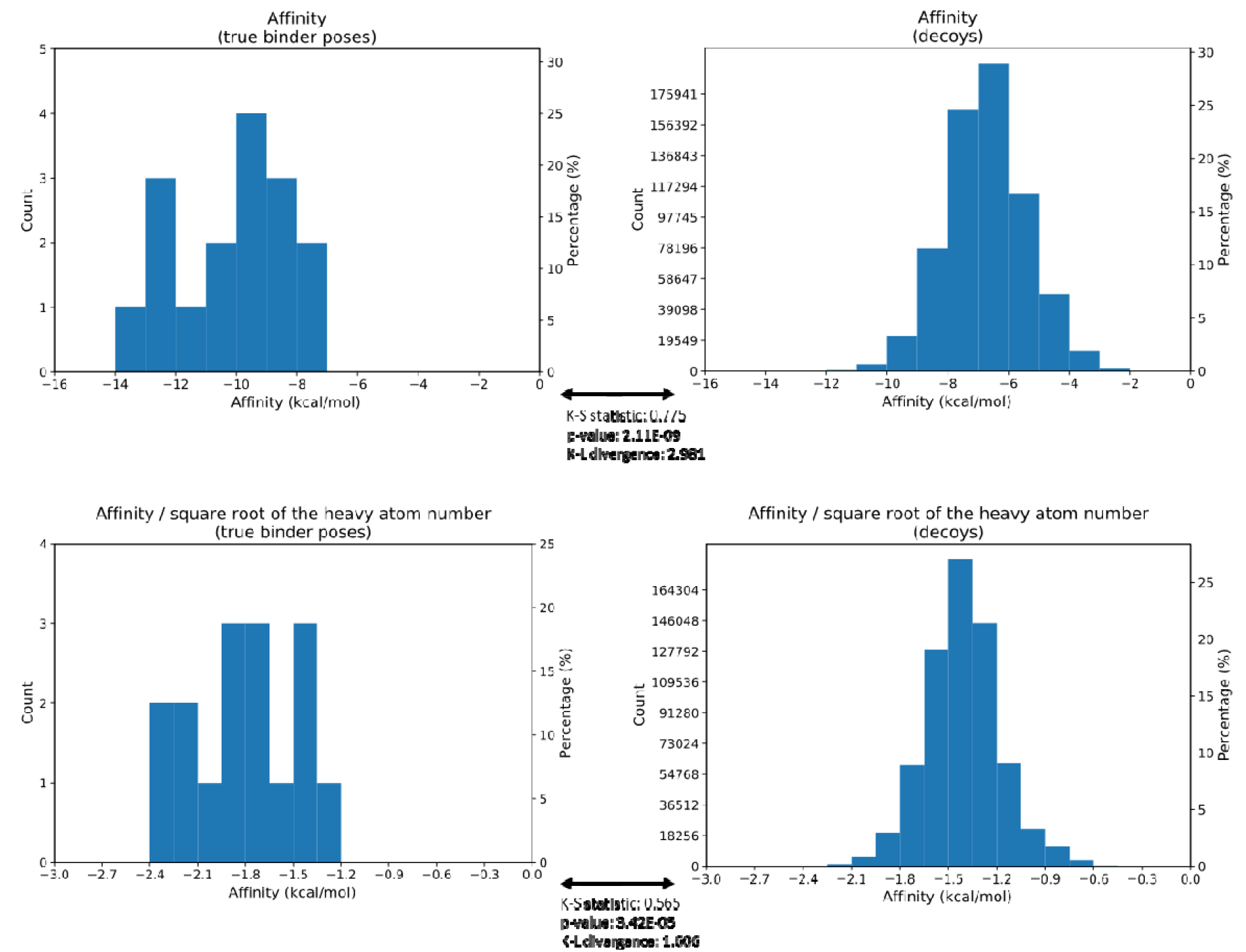
Difference in the distributions of the affinities between the “answer” poses (the true binder pose, left columns) and decoys (right columns) assessed by two different metric - Kolmogorov–Smirnov (K-S) test and Kullback–Leibler (K-L) divergence. Figures shows the original Vina predicted affinity values (top row) or normalized affinity by the square root of drug heavy atom number (bottom row). K-S statistic, the statistic in the Kolmogorov–Smirnov test, which tests against the null hypothesis that there is no difference between the distributions. p-value: the p-value in the distribution of the K-S statistic. The smaller the value, the more significant the difference. K-L divergence: Kullback–Leibler divergence, which measures the difference between the two distributions. The larger the value, the more different the two distributions. The results showed that the answer poses and decoys have more different distributions on the original affinity values than the normalized affinity values.

**Figure 3.**
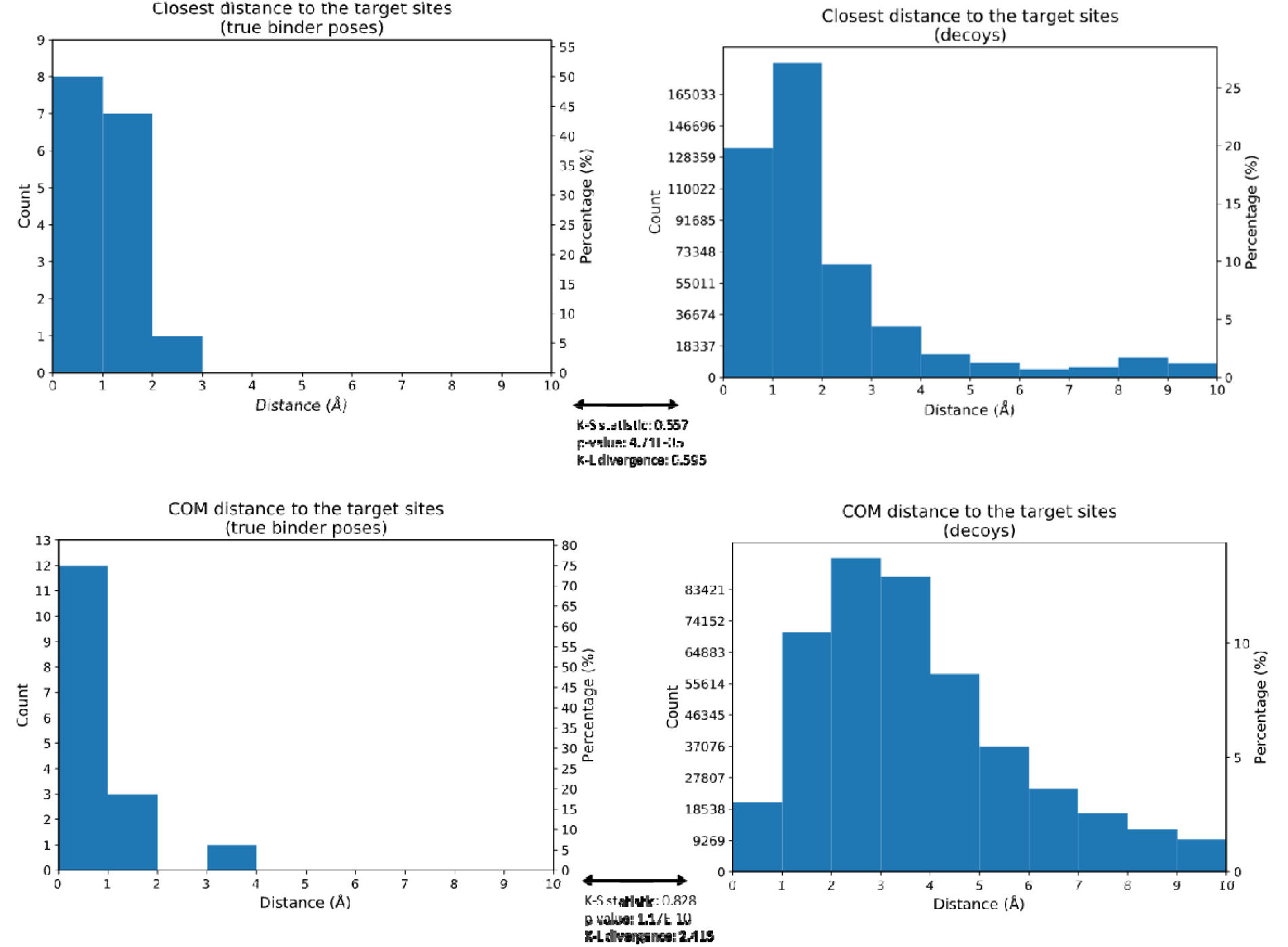
The comparison of the difference in the distribution of the distance to the target sites for the answer poses (left columns) and decoys (right columns) measured by the closest distance between any heavy atoms (top row) or by the distance from the center of mass (COM) of the drug. K-S statistic, the statistic in the Kolmogorov–Smirnov test. p-value: the p-value in the distribution of the K-S statistic. K-L divergence: Kullback–Leibler divergence. The results showed that the answer poses and decoys have more different distributions on using the closest distance as the distance measure than using COM distance.

**Figure 4.**
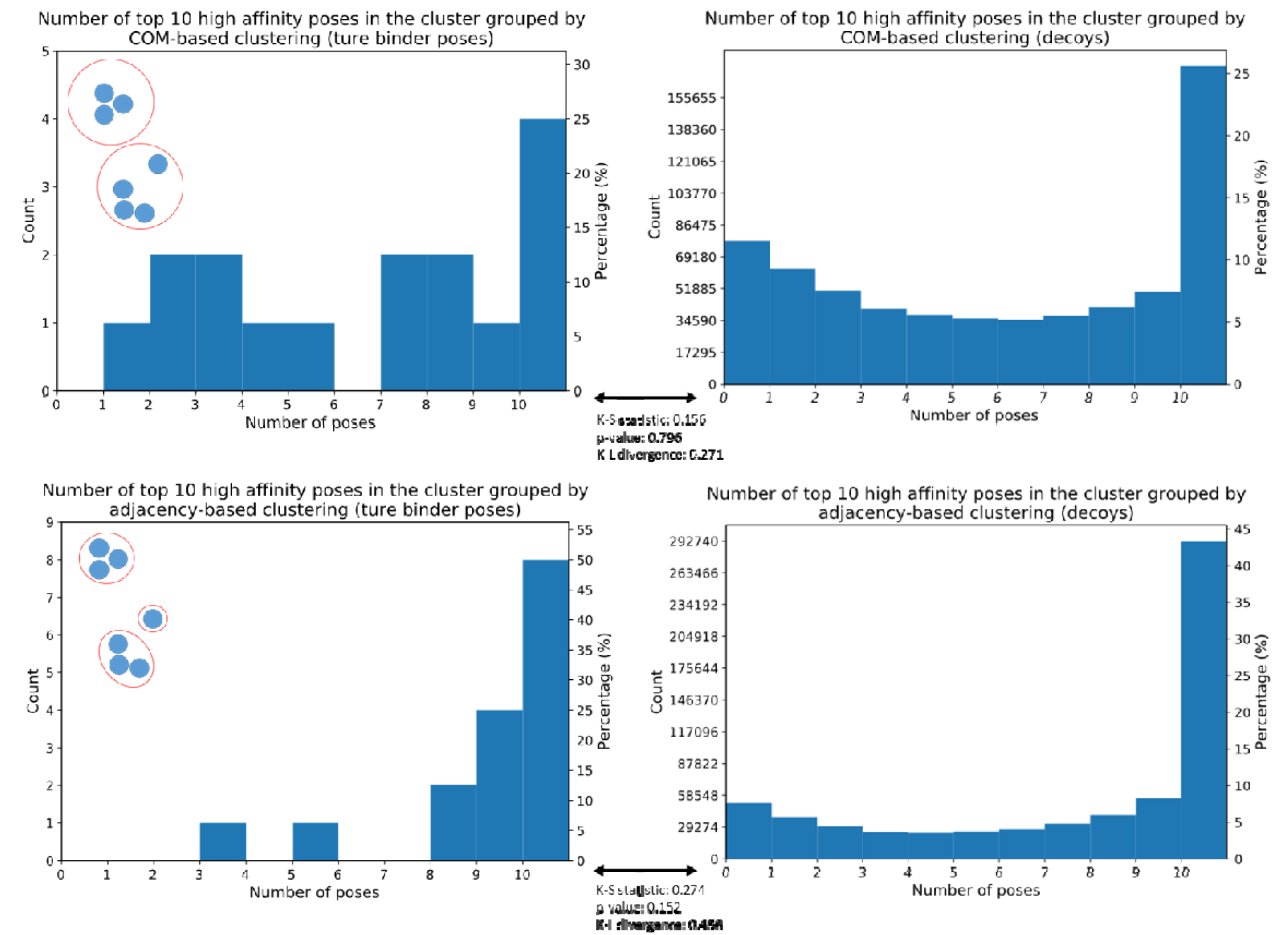
The comparison of the difference in the distribution of the number of high-affinity poses (cluster size), defined as the top 10 highest affinity poses, in the cluster grouped by COM-based clustering (top row) or adjacency-based clustering (bottom row) for the answer poses (left columns) and decoys (right columns). K-S statistic, the statistic in the Kolmogorov–Smirnov test. p-value: the p-value in the distribution of the K-S statistic. K-L divergence: Kullback–Leibler divergence. The results showed that the answer poses and decoys have more different distributions on the number of high-affinity poses in the clustered grouped by adjacency-based clustering than COM-based clustering.

All the three drug ranking methods proceeded in two stages-a representative pose among the sampled poses was first chosen for each drug and the drugs were then ranked based on their representative poses. The performance of a method can be evaluated using two metrics - (A) the averaged ranking of true binders (Eq. 4) as well as (B) the percentage of true binders being prioritized to the top 5% or 10% of the examined drugs (Tables 2 and 3). To dissect the contribution of individual features, we computed the performance of the *ad hoc* method using only affinity, both affinity and distance, and all of the three features. The ranking based on affinity showed an average ranking of 164.44 and with 75% of the true binder drugs ranking in the top 10% of the 2016 FDA drugs (Table 2). When taking the distance and cluster size into account, the ranking was slightly improved and resulted in an average rank of 153.81 and that 81.3% of the true binders are prioritized to the top 10%, suggesting the usefulness of adding the distance and cluster features. However, when looking at the Test Set results (Table 3), we found using all three features do not benefit the *ad hoc* results but using all three or just the “affinity + distance” benefit the FRS. Overall, the drug ranking predicted by the LOD scores suffer the least bias from the special case (see the results for 4zau) and showed a significant improvement in the average rank of the co-crystallized drugs in both training (Avg. rank: 110.5, Table 2) and testing dataset (Avg. rank: 26.75, while other methods rank >232, Table 3), which also reflected on the improved success rate of 100% (Table 3). The *ad hoc* method adopted a hierarchical strategy to rank features in different order, which could introduce biases. For a “loser” in the beginning hardly overcome the disadvantage for other outstanding features. On the contrary, the LOD score considered the three features altogether, and drugs getting a high score, suggesting relatively lower probability in the decoy set to have that feature value, in any of the three features were then possible to stood out and thus reducing the aforementioned problem.

**Table 2.**
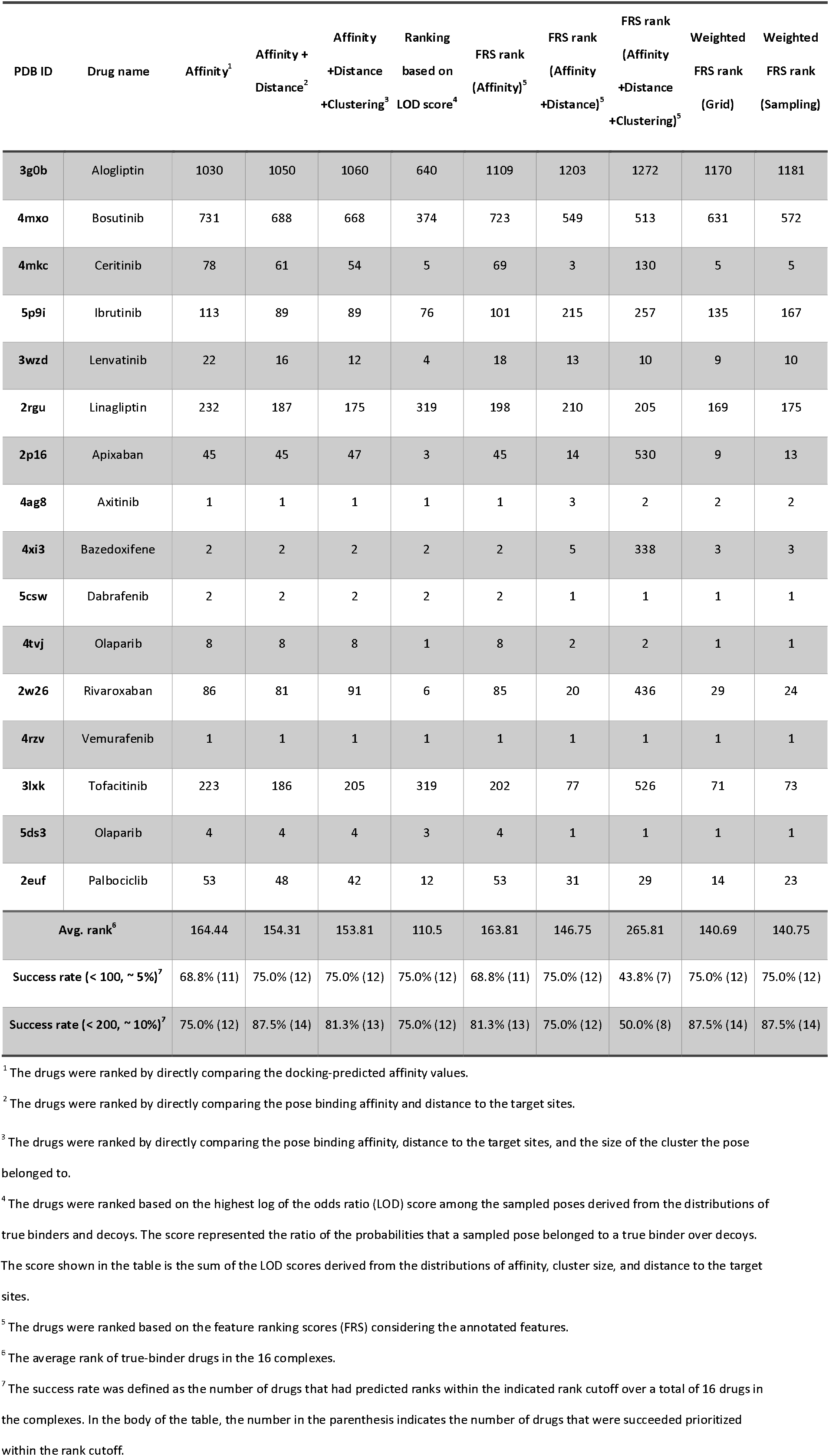
The predicted ranking of the answer drugs, the FDA-approved drugs bound in our collected 16 complexes, among the screened 2016 drugs, the average ranking, and the success rates by different ranking methods. The success rate was defined as the proportion of drugs in the 16 complexes that were ranked within the indicated top rank 100 (∼5%) or 200 (∼10%).

**Table 3.** The predicted ranking of the answer drugs for the four protein targets in the testing set.

| PDB ID | Drug name | Affinity<sup>1</sup> | Affinity + Distance<sup>2</sup> | Affinity + Distance + Clustering<sup>3</sup> | Ranking based on LOD score<sup>4</sup> | FRS rank (Affinity)<sup>5</sup> | FRS rank (Affinity + Distance)<sup>5</sup> | FRS rank (Affinity + Distance + Clustering)<sup>5</sup> | Weighted FRS rank (Grid) | Weighted FRS rank (Sampling) |
| --- | --- | --- | --- | --- | --- | --- | --- | --- | --- | --- |
| 2wgj | Crizotinib | 41 | 38 | 39 | 2 | 41 | 1 | 339 | 1 | 1 |
| 6nec | Nintedanib | 21 | 20 | 17 | 3 | 21 | 1 | 1 | 1 | 1 |
| 4zau | Osimertinib | 585 | 738 | 712 | 86 | 585 | 470 | 301 | 739 | 650 |
| 4bjc | Rucaparib | 284 | 249 | 244 | 16 | 284 | 6 | 6 | 21 | 16 |
| Avg. rank<sup>6</sup> |  | 232.75 | 261.25 | 253.0 | 26.75 | 232.75 | 119.5 | 161.75 | 190.5 | 167.0 |
| Success rate (< 100, ~ 5%)<sup>7</sup> |  | 50.0% (2) | 50.0% (2) | 50.0% (2) | 100.0% (4) | 50.0% (2) | 75.0% (3) | 50.0% (2) | 75.0% (3) | 75.0% (3) |
| Success rate (< 200, ~ 10%)<sup>7</sup> |  | 50.0% (2) | 50.0% (2) | 50.0% (2) | 100.0% (4) | 50.0% (2) | 75.0% (3) | 50.0% (2) | 75.0% (3) | 75.0% (3) |

<sup>1, 2, 3, 4, 5, 6, 7</sup> The definitions are the same as that in Table 2 and the average ranking and success rates by different ranking methods are also defined in the same way as Table 2.

For the FRS, our results showed that FRS performed the best when considering both the binding affinity and the distance to the active site, with an average ranking of 146.75 in the training set and 119.5 in the test set, both having 75% of the top 10% success rate (Table 2). We then compared the performance of the weighted FRS scores refined by the grid-search and the sampling method, which gave the weights of (α, β, γ) = (0.8, 0.2, 0) and (0.72470771, 0.27438212, 0.00091017), where α, β, and γ were the weights for the binding affinity, distance to the active site, and cluster size, correspondingly. The learned weights suggested that the cluster size failed to prioritize these co-crystallized drugs by this method, which was consistent with the poor performance of FRS when considering all the three features (with equal weights) (Tables 2 and 3). The average rank of the weighted FRS (Avg. rank: 140.69 and 140.75) outperformed the equal weights FRS (Avg. rank: 163 to 266) in the training dataset, but did not show as competitive performance (Avg. rank: 190.5 and 167.0) compared to the equal weights FRS using affinity and distance features (Avg. rank: 119.5) in the testing dataset. *Our results suggested that the LOD score and the FRS based on affinity and distance features giving the best outcome to prioritize the co-crystallized drugs among the decoys*, from small molecule docking. Given that its best and consistent performance in both the training and testing dataset, the LOD score was chosen as the ranking method embedded in our DRDOCK.

## 5 Application

Coronavirus disease 2019 (COVID-19), an outbreak that began by the end of 2019, had progressed to a global pandemic in 2020 and severely impacted human society in whole aspects. Efforts for finding effective and safe treatment besides vaccines are still being made. In the host cell, the severe acute respiratory syndrome (SARS) coronaviruses, including SARS-CoV-2, the causality of COVID-19, methylated the 5′ cap structure of newly synthesized mRNAs to mimic the native host mRNAs and prevent their recognition and degradation by the host immune system (Daffis et al., 2012). The virus non-structural protein 16 (nsp16), which formed a complex with nsp10 (nsp16/10), was responsible for this methylation by transfer the methyl group from the methyl donor, S-adenosyl methionine (SAM), to the first nucleotide of the mRNA at 2′-O of the ribose sugar (Krafcikova et al., 2020; Viswanathan et al., 2020). The nsp16, a cap ribose 2′-O methyltransferase, thus became a potential drug target where peptides or drugs that could inhibit its enzyme activity could raise the instability of viral mRNAs and interferon (IFN)-mediated antiviral response (Wang et al., 2015), and, therefore, apply to COVID-19 treatment. Here, we used our webserver to screen drugs as the inhibitors for SARS-CoV2 nsp16. We used the nsp16 structure from SARS-CoV2 (PDB Code: 6wks, chain A) as the protein target submitted to our webserver. The position of the natural ligand, SAM (residue ID: 301), was chosen as the target site, of which the COM was used to calculate the distance from the docking poses that was used in LOD scores calculation and drug ranking. We then further submitted the top-ranked 20 drugs from the docking results to 10 ns MD simulations and got the corresponding MM/GBSA energies from the sampled snapshots.

Table 4 showed the top-ranked 20 drugs based on the LOD scores derived from the docking results and their corresponding indicators inferred from 10 ns MD simulations, including the mean binding free energy, the drug leaving time, and the largest distance of the drug to the target sites during the simulations (Methods 2.9). The natural ligand, S-adenosyl methionine (SAM), serving as a methyl donor for the mRNA substrate and belonging to one of the FDA-approved drugs included in our drug library, was ranked 17 at the docking stage, which demonstrated the robustness and accuracy of our drug ranking method based on the LOD score (Methods 2.6). Confirmed by the 10 ns simulation, SAM was further prioritized to rank 2 with the mean binding free energy of -31.0 kcal/mol and the longest distance of 1.43 Å to the target site, suggesting the high affinity and stable binding of the natural ligand with nsp16. Among the top 20 ranked drugs, several drugs failed to form tight interactions with the SAM binding pocket when examined by MD and finally left the binding pocket (distance > 10 Å from the target site) during the 10 ns simulations, including olaparib (docking rank: 2, MD rank: 15, drug leaving time: 6.9 ns), levosimendan (docking rank: 3, MD rank: 19, drug leaving time: 1.3 ns), and ataluren (docking rank: 4, MD rank: 17, drug leaving time: 4.2 ns), which suggested the limitation of docking scoring function with a fixed receptor at zero temperature (no kinetics) in vacuum. The re-evaluation of the binding by MD simulations in explicit-solvent could benefit the overall drug screening outcome.

**Table 4.** The top-ranked 20 drugs by the LOD score from nsp16’s docking results.

| Drug name | Docking rank<sup>1</sup> | LOD score<sup>2</sup> | Affinity (kcal/mol)<sup>3</sup> | Affinity rank<sup>4</sup> | MD rank<sup>5</sup> | Mean binding free energy (kcal/mol)<sup>6</sup> | Drug leaving time (ns)<sup>7</sup> | Longest distance to the target site (Å)<sup>8</sup> |
| --- | --- | --- | --- | --- | --- | --- | --- | --- |
| Enasidenib | 1 | 5.31 | -8.8 | 104 | 7 | -18.64 | - | 2.97 |
| Olaparib | 2 | 5.13 | -9.9 | 16 | 15 | -20.86 | 6.9 | 12.18 |
| Levosimendan | 3 | 5.03 | -7.9 | 378 | 19 | -5.69 | 1.3 | 29.56 |
| Ataluren | 4 | 5.03 | -7.9 | 379 | 17 | -10.54 | 4.2 | 24.48 |
| Nifuroxazide | 5 | 5.03 | -7.5 | 583 | 10 | -16.46 | - | 1.75 |
| Fenoterol | 6 | 5.03 | -7.4 | 646 | 4 | -24.41 | - | 6.64 |
| Duvelisib | 7 | 4.6 | -9.1 | 69 | 16 | -14.58 | 5 | 10.41 |
| Carprofen | 8 | 4.49 | -7.7 | 460 | 18 | -6.41 | 2.1 | 22.85 |
| Ergotamine | 9 | 4.12 | -10.5 | 4 | 8 | -17.53 | - | 5.41 |
| Sm13496 | 10 | 4.12 | -10.4 | 6 | 5 | -24.17 | - | 5.22 |
| Raltegravir | 11 | 3.73 | -9.8 | 18 | 14 | -11.62 | 9.2 | 14.09 |
| Abemaciclib | 12 | 3.73 | -9.3 | 46 | 9 | -16.68 | - | 5.21 |
| Fluralaner | 13 | 3.73 | -9.1 | 70 | 1 | -35.56 | - | 3.33 |
| Olmesartan | 14 | 3.73 | -9.1 | 71 | 20 | -13.02 | 0.8 | 13.28 |
| Dantrium | 15 | 3.63 | -7.9 | 380 | 12 | -12.18 | - | 9.55 |
| Tafamidis | 16 | 3.63 | -7.6 | 514 | 11 | -13.38 | - | 7.74 |
| S-adenosyl-methionine | 17 | 3.63 | -7.6 | 515 | 2 | -31.00 | - | 1.43 |
| Tegaserod | 18 | 3.63 | -7.2 | 828 | 3 | -25.34 | - | 3.47 |
| Conivaptan | 19 | 3.26 | -10.6 | 3 | 6 | -19.14 | - | 6.92 |
| Eltrombopag | 20 | 3.26 | -10.3 | 7 | 13 | -26.2 | 9.3 | 11.32 |

<sup>1</sup> The rank among 2016 drugs based on the log of the odds ratio (LOD) score (column 2).
<sup>2</sup> The log of the odds ratio (LOD) score used to rank the drugs based on the docking results. See Method 3.5.
<sup>3</sup> Docking affinity predicted by AutoDock Vina.
<sup>4</sup> The rank among 2016 drugs based on the docking affinity (column 4).
<sup>5</sup> The rank among the top 20 drugs subject to further 10 ns simulations. The drugs were ranked based on the average binding free energies (column 7) and drug leaving times (column 8).
<sup>6</sup> The average binding free energy by MM/GBSA method (Kollman et al., 2000; Massova et al., 2000) over the sampled snapshots during the simulations.
<sup>7</sup> The simulation time when the drug dissociated from the target site (the first time point when the distance of the drug to the target site was > 10 Å). -, the drugs did not dissociate during the whole 10 ns simulations.
<sup>8</sup> The longest distance of the drug to the target site sampled during the 10 ns simulations.

Our webserver also suggested several drugs that could potently bind and compete with SAM for its binding pocket, as reflected on their top-ranked LOD scores, favorable mean MM/GBSA binding free energies, and tight adhesion in the binding pocket suggested by the small departure of the drugs from the target site (“longest distance during simulation to the target site”) (Methods 2.9). By analyzing the MM/GBSA-revealde binding affinity from MD trajectories, we took top four drugs except for the native substrate SAM – Fluralaner (−35.56 kcal/mol), Tegaserod (−25.34 kcal/mol), and Fenoterol (−24.41 kcal/mol). All three cannot be found in the top 5 of the original docking results with Fluralaner and Tegaserod ranking 13 and 18, respectively, where 2 out of 3 could stably stay within 4 Å from the target site (Table 4).

We further test how these three drugs suppress virus replication in SARS-CoV2-infected Vero E6 cells (hCoV-19/Taiwan/NTU13/2020; Accession ID: EPI_ISL_422415). The plaque reduction by the drug repurposing can be found in Figure 5. Fluralaner is an insecticide used to treat fleas and ticks in animals and recently proposed to treat vector-borne diseases including malaria and Zika fever (Miglianico et al., 2018). It was found toxic to Vero E6 cells at 10 μM and therefore the antiviral effect could not be shown. Yet, we delightfully report here that Tegaserod and Fenoterol exhibit anti-SARS-CoV2 activities at 10 and 40 μM respectively, reducing the plaques by 54% and 70%. Tegaserod activating the 5-HT4 receptors in the enteric nervous system is formulated in tablet for oral delivery to treat irritable bowel syndrome with constipation (IBS-C) (Camilleri, 2001; Black et al., 2020). On the other hand, Fenoterol is a β-adrenoreceptor agonist (Svedmyr, 1985) and currently sold as an inhaled bronchodilator for an asthma attack. Asthma has been associated with worsening symptoms in patients and is generally considered a risk factor of COVID-19 severity (Johnston, 2020). Centers for Disease Control and Prevention of American also reminds that asthma may cause severer illness from COVID-19 infection and those patients with asthma should take more care and preventive actions (Centers for Disease Control and Prevention, 2021). The moderate suppression of SARS-CoV2 suggested that Fenoterol could be a timely reliever for COVID-19 patients with asthma.

**Figure 5.**
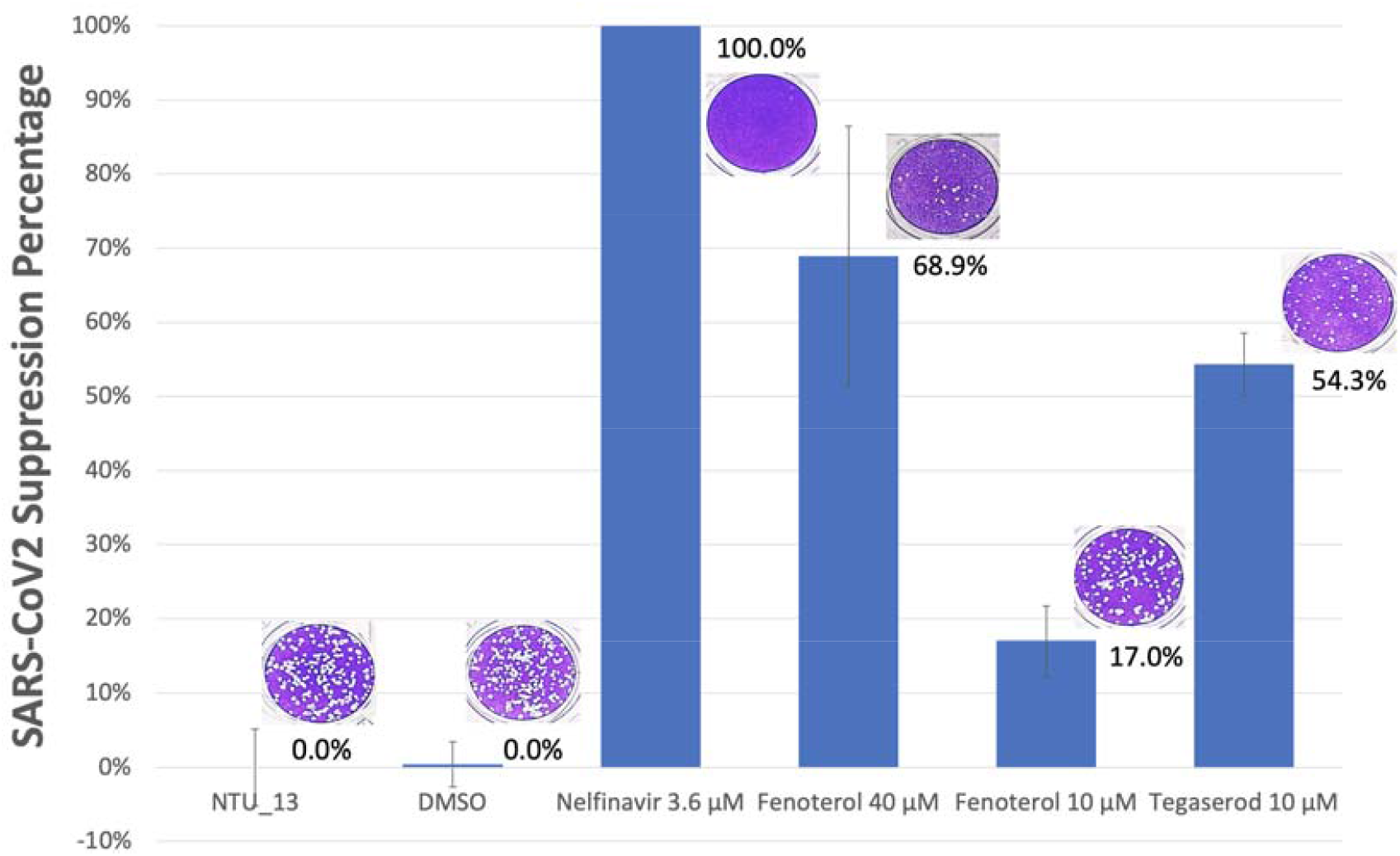
**Plaque reduction percentage of two repurposed SARS-CoV2 nsp16 active site inhibitors tegaserod and fenoterol in comparison with an earlier reported SARS-CoV2 3CL protease inhibitor, Nelfinavir** that serves as a positive control (Pathak et al., 2021) in this experiment. The inhibition percentage is calculated as [1-(V_D_/V_C_)]×100%, where V_D_ and V_C_ refer to the virus titer in the presence and absence of the inhibitors, respectively. DMSO indicates DMSO/buffer mix where the DMSO% is less than 0.8% as a mock drug and negative control. In this experiment, drug free medium and the DMSO control happen to result in the same average number of plaques. Representative plaque phenotype for each group is placed on top of the bar. Vero E6 cells are stained by crystal violet and white spots correspond to virus-infected and -burst cells.

We hope our platform provides a timely suggestion for drug benchmarking and re-ranking as well as a convenient tool (fueled with heavy computing in the cloud end) for drug repurposing used in current and future outbreak of emergent diseases. Our rationales of making available this service are further discussed in the Supporting Information.

