## Supplemental Methods for "DRDOCK: A Drug Repurposing platform integrating automated docking, simulations and a log-odds-based drug ranking scheme"

**Supplementary Information**

**Contents**

**1. Supplementary methods**

**Benchmark set curated for the development of drug ranking methods**

**2. Supplementary results and discussion**

**DRDOCK – user submission**

**3. Supplementary tables**

**Table S1** Supporting information of the collected 20 PDB complexes containing FDA-approved drugs used in this study.

**4. Supplementary figures**

**Figure S1** *Ad hoc* method for drug ranking.

**Figure S2** The user submission form and results page of DRDOCK.

**5. References**

**1. Supplementary methods**

**Benchmark set curated for the development of drug ranking methods**

To develop the drug ranking methods and compare their performance, the complex structures for a set of known target proteins and FDA-approved drugs were extracted from the Protein Data Bank (PDB) (rcsb.org; Berman et al., 2000). We first collected 48 chemical IDs of FDA drugs that were approved during 2010-2016 gathered in the article published by Westbrook et al (2019) (the drugs reported to target the same target proteins were excluded for simplicity). Among these 48 approved drugs, 44 were included in our 2016 FDA-approved drug library. We then fetched the PDB IDs in which the structures were co-crystallized with one of the 44 drugs. Those PDB IDs of the drug targets that were the same as the targets of the complexed drugs annotated in Westbrook’s article (by matching the UniProt IDs) were retained. For those drugs of which any of the complexed targets of the found PDB IDs were matched to the annotated target, we compared their Enzyme Commission number (EC number) and kept those PDB IDs that shared the same EC number. This resulted in 86 PDB IDs composed of 26 drug targets for 33 out of the aforementioned 44 drugs. From these, we retained those protein structures solved by X-ray crystallography, without nonnative amino acids, and from Homo sapiens. We also eliminated those housing the drug binding sites occupied by ions and/or co-crystallizers that contact the FDA drugs of interest (within 5 Å), those containing missing residues within 5 Å of the bound drug that could impair the integrity of the binding site, those with documented low affinities (Kd, Ki, IC50 > 1000 nM), and those belonging to the membrane protein. This resulted in 40 PDB IDs composed of 24 non-redundant target protein-FDA drug pairs (10 targets have more than 2 PDB IDs) for 18 target proteins and 19 FDA drugs. For each target protein-FDA drug pair that had several PDB IDs, we followed the ordered criteria to choose one PDB ID as the representative: the target protein did not have mutated residues, having annotated binding affinity, having a higher resolution, and having higher binding affinity. This resulted in 20 non-redundant protein-drug complexes among which 16 were randomly assigned as the training set and the remaining four as the testing set (Table S1). We called the FDA-approved drug in the complex structure as “true binder”.

**2. Supplementary results and discussions**

**DRDOCK – user submission**

DRDOCK requires users to submit a well-prepared single-chain protein structure in PDB format and the corresponding residues IDs of the target sites, such as active sites for enzymes, that are used to identify the closest docking poses the users should interest in (Figure S2A). Since the integrity of the protein structure is essential for meaningful MD simulations, the protein structure should be prepared for preventing any missing backbone heavy atoms (CA, C, N, O). The missing of any side-chain heavy atoms is allowed since they can be automatically patched according to pre-defined amino acid templates but may slightly distort the predicted binding poses in docking. DRDOCK requires the protein structure to only comprise 20 standard amino acids, of which the force constants and atom charges are already well developed and calibrated in AMBER ff14SB force field (Maier et al., 2015). For those structures containing some of the standard amino acid-derived residues, DRDOCK can tolerate that by mutating the residues back to their closest standard amino acids. Otherwise, the users should manually modify and mutate back those residues according to the original protein sequence. The most convenient way to achieve this is to submit the desired target protein structure with non-standard amino acids with its corresponding original peptide sequence to the SWISS-MODEL web server (Waterhouse et al., 2018) under the “template mode”, which can automatically mutate any residue that is different from the original peptide sequence back and patch any missing heavy atoms, which is our recommended way to prepare a submission-ready target structure.

**Discussions**

Drug repurposing can usually serve as the first possible remedy in any emergent life-threatening infectious disease, before vaccines and any drug with high specificity can be made available. To promptly identify effective drugs from existing ones, a drug screening platform with both efficiency and accuracy is required (Park, 2019; Pulley et al., 2020; Pushpakom et al., 2018). Our web server aims to fulfill this prospect by providing an automatic drug screening platform for drug repurposing of 2016 FDA-approved drugs toward any protein target. Its easy-to-use interface removes the technical barrier of virtual screening and helps accelerate drug repurposing for abrupt outbreaks of merging infectious diseases.

A general goal for virtual screening is to enrich the effective drugs for a given protein target to the top of a set of decoys (Graves et al., 2006) in a drug library. It is, therefore, more rationalized to use a scoring function trained based on both the true binders and decoys. However, there are difficulties when trying to include decoys in the calibration process. First, the affinity of a decoy is too weak to be experimentally measured. Second, the atomic interaction between the protein and a decoy is not available without a solved complex structure. This results in that most scoring functions are calibrated using known protein-ligand complex structures with experimentally measured affinity while not taking decoys into account, although the evaluation of a ligand affinity based on the known atomic interaction is thought to be generally applied to either true binders and decoys. Here, we proposed a compromised way where we call a drug known to bind a protein target as the true binder and other drugs in the drug library as decoys for that target. A successful ranking method should prioritize the true binder on top of the decoys as much as possible. With this rationale, we defined a loss function as the total sum (or average if divided by the number of targets) of the rankings assigned by a ranking method for the true binders in the collected set of known protein-drug complexes. This loss function can be used to evaluate existed or search for improved ranking methods and is generally applicable to any set of protein-drug complexes.

To learn from both true binders and decoys, the atomic details of the proteins-drugs interaction are generated by docking software, Vina. The sampled docking poses form the statistical basis for us to distinguish the true binders from the decoys based on the different distributions of their pose features. The most distinguishable features include the affinity from Vina, the distance from pose COM to the target site, and the size of a pose cluster that inversely reflected the degree of entropy loss during the drug binding. A log-odds (LOD) score is designed as a quantitative tendency for describing a sampled pose is more likely true binders or decoys. We also propose a feature ranking score (FRS) that aims to avoid systematic bias when applying the method to different protein targets. Our results show that the LOD score had an outstanding improvement on the average ranking of the true binders consistent in both training and testing datasets, suggesting its general applicability. This also demonstrates that the incorporation of the decoys properties do help improve the correctness of the drug ranking methods.

While docking allows the screening of a huge drug library within a feasible time, the accuracy of predicted ligand binding affinity is hindered by its simplified scoring function, evaluating in a vacuum environment instead of water, and not taking the dynamics of protein-drug interaction into account (Pantsar et al., 2018). This is addressed by MD simulations with delicate force fields in explicit solvent. In DRDOCK, the pre-built parameters files of 2016 FDA-approved drugs and a straightforward user interface allow advanced binding free energy evaluation by MD simulations in just a few clicks. The double confirmations reduce the top-ranked decoys that are simply wasted if subject to experimental assays and shall help raise the success rate in drug repurposing.

**3. Supplementary tables**

**Table S1**. Supporting information of the collected 20 PDB complexes containing FDA-approved drugs used in this study.

| PDB ID | UniProtKB ID | Target Name | Enzyme | Classification | Drug target site^1^ | References |
| --- | --- | --- | --- | --- | --- | --- |
| Training Set | | | | | | |
| 3g0b | P27487 | Dipeptidyl peptidase 4 | Yes | Serine protease | Catalytic site | Zhang et al., 2011 |
| 4mxo | P12931 | Proto-oncogene tyrosine-protein kinase Src | Yes | Non-receptor tyrosine kinase | ATP binding site | Levinson et al., 2013 |
| 4mkc | Q9UM73 | ALK tyrosine kinase receptor | Yes | Receptor tyrosine kinase | ATP binding site | Friboulet et al., 2014 |
| 5p9i | Q06187 | Tyrosine-protein kinase BTK | Yes | Non-receptor tyrosine kinase | ATP binding site | Bender et al., 2017 |
| 3wzd | P35968 | Vascular endothelial growth factor receptor 2 | Yes | Receptor tyrosine kinase | ATP binding site | Okamoto et al., 2014 |
| 2rgu | P27487 | Dipeptidyl peptidase 4 | Yes | Serine protease | Catalytic site | Eckhardt et al., 2007 |
| 2p16 | P00742 | Coagulation factor X | Yes | Serine protease | Catalytic site | Pinto et al., 2007 |
| 4ag8 | P35968 | Vascular endothelial growth factor receptor 2 | Yes | Receptor tyrosine kinase | ATP binding site | Mctigue et al., 2012 |
| 4xi3 | P03372 | Estrogen receptor | No | Nuclear receptors | Hormone binding site | Fanning et al., 2016 |
| 5csw | P15056 | Serine/threonine-protein kinase B-raf | Yes | Serine/threonine protein kinase | ATP binding site | Waizenegger et al., 2016 |
| 4tvj | Q9UGN5 | Poly [ADP-ribose] polymerase 2 | Yes | Poly (ADP-ribose) polymerase | Nicotinamide pocket | Thorsell et al., 2016 |
| 2w26 | P00742 | Coagulation factor X | Yes | Serine protease | Catalytic site | Roehrig et al., 2005 |
| 4rzv | P15056 | Serine/threonine-protein kinase B-raf | Yes | Serine/threonine protein kinase | ATP binding site | Karoulia et al., 2016 |
| 3lxk | P52333 | Tyrosine-protein kinase JAK3 | Yes | Non-receptor tyrosine kinase | ATP binding site | Chrencik et al., 2010 |
| 5ds3 | P09874 | Poly [ADP-ribose] polymerase 1 | Yes | Poly (ADP-ribose) polymerase | Nicotinamide pocket | Dawicki-Mckenna et al., 2015 |
| 2euf | Q01043 | Cyclin homolog | Yes | Cyclin-dependent kinase | ATP binding site | Lu et al., 2006 |
| Test Set | | | | | | |
| 2wgj | P08581 | Hepatocyte growth factor receptor | Yes | Receptor tyrosine kinase | ATP binding site | Cui et al., 2011 |
| 6nec | P07949 | Proto-oncogene tyrosine-protein kinase receptor | Yes | Receptor tyrosine kinase | ATP binding site | Terzyan et al., 2019 |
| 4zau | P00533 | Epidermal growth factor receptor | Yes | Receptor tyrosine kinase | ATP binding site | Yosaatmadja et al., 2015 |
| 4bjc | Q9H2K2 | Poly [ADP-ribose] polymerase tankyrase-2 | Yes | Poly (ADP-ribose) polymerase | Nicotinamide pocket | Haikarainen et al., 2013 |

^1^ The functional site targeted by the drug.

**4. Supplementary figures**

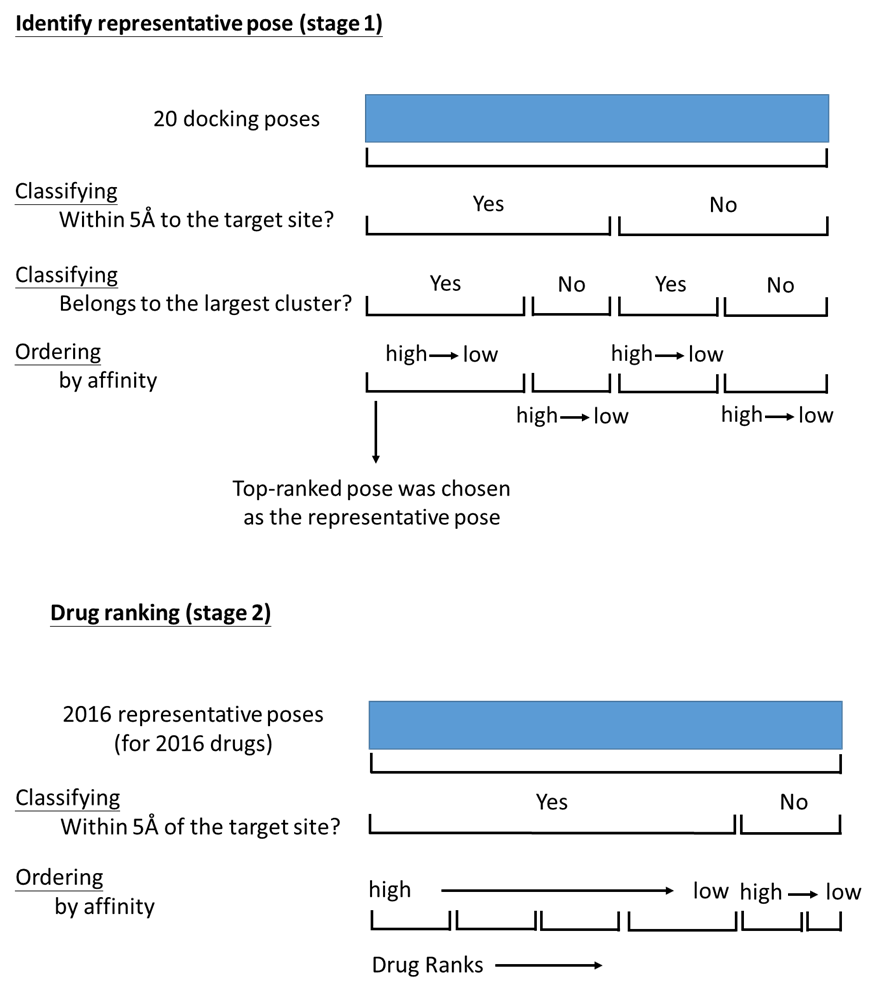

**Figure S1**. ***Ad hoc* method for drug ranking**. The poses and drugs were ordered in a hierarchical order-by-group manner in two stages. In the first stage, the sampled 20 poses for each drug were ordered and a representative pose was chosen as the top-ranked pose. This included two steps of hierarchical binary classifications, based on features of the pose’s distance to the target site and the size of the pose cluster, and resulted in four groups of poses. Within each group, the poses were then sorted based on the pose affinity. In the second stage, the rankings of 2016 drugs were determined using the representative poses chosen in the first stage. The drugs were first classified and grouped into two based on whether the docking pose was within 5 Å of the target site. The final drug rankings were then determined by ordering the drugs from high to low affinity in each of the groups.

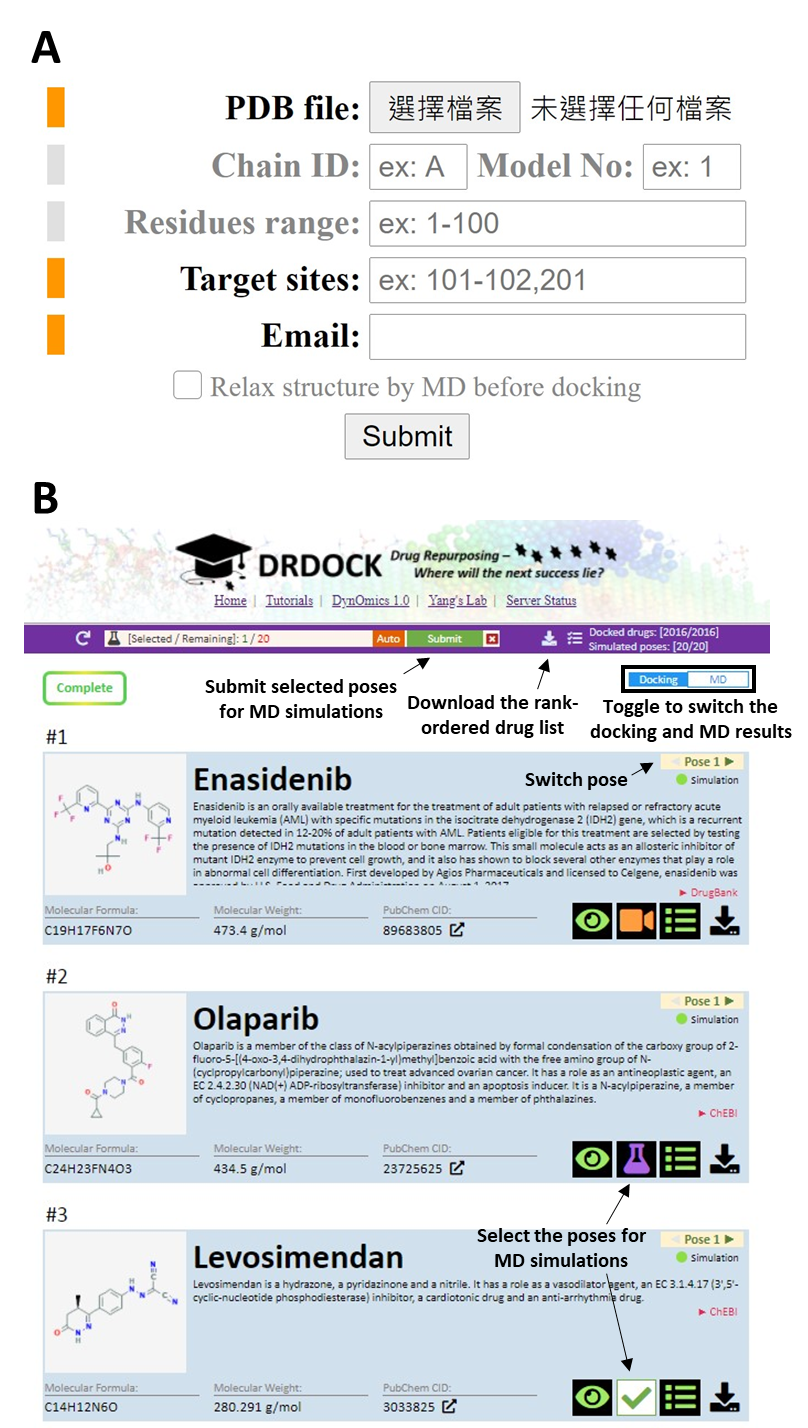

**Figure S2**. The user submission form and results page of DRDOCK. **(A)** The job submission form in the main page of DRDOCK. The user is required to provide a well-prepared single-chain protein structure (“PDB file”). The user can assign a specific chain ID, model number for NMR-resolved structures, a desired segment in the specified protein chain (in “Residues range”). Users are required to specify the IDs of residues that comprise a target site (e.g. enzyme active site residues). Lastly, the user should provide a valid email address for receiving the job status and results from DRDOCK. The orange fields are required while grey fields are optional. **(B)** The results page of DRDOCK. The drugs are rank-ordered based on our ranking algorithm that ranks docked poses as described in the Methods. The drug name, 2D structure, description, basic properties, and PubChem link are shown in the drug panel. The rank-ordered drug list in CSV format can be downloaded (the download button in the purple bar atop). To submit poses for MD simulations, click the flask icon to enter the simulation submission mode, where a submission panel and a “submit” button will show up in the functional bar atop (beneath the navigation bar). A checkmark also shows up and replaces the flask icon for chosen drugs. Users can also switch the docking poses to submit different poses for the same drug. Finally, click the “submit” button in the submission panel to submit all the checked poses to the web server. Users can browse the rank-ordered drugs based on the simulation results (see Methods) by toggling the “MD” at the upper right corner (highlighted in the black box in **Fig S2B**). See more functions in Figure 1C for visualizing the docking pose and playing the trajectory. Other details about DRDOCK, including that our file parser can allow several types of non-standard amino acids, can be found in **Supplementary methods**.
